# Rectal bacteria produce sex pheromones in the male oriental fruit fly

**DOI:** 10.1101/2020.12.17.423356

**Authors:** Lu Ren, Ma Yingao, Mingxue Xie, Yongyue Lu, Daifeng Cheng

## Abstract

In recent decades, a growing body of literature has indicated that microbial symbionts of insects can modulate their hosts’ chemical profiles and mate choice decisions. However, there is currently little direct evidence indicate that insect pheromones can be produced by symbionts. Using *Bactrocera dorsalis* as a model system, we demonstrate that *Bacillus sp.* in the rectum of male *B. dorsalis* plays a pivotal role in sex pheromones production. We demonstrate that 2,3,5-trimethylpyrazine (TMP) and 2,3,5,6-tetramethylpyrazine (TTMP) are sex pheromones produced in the male rectums. Mature virgin females can be strongly attracted by TMP and TTMP. TMP and TTMP contents in male rectums can be decreased when rectal bacteria are inhibited with antibiotics. Moreover, *Bacillus* sp. isolated from male rectum can produce TMP and TTMP when providing with substrates-glucose and threonine, for which the contents are significantly higher in rectums of mature males. These findings highlight the influence of microbial symbionts on insect pheromones and provide an example of direct bacterial production of pheromones in insects.

## Introduction

In a dynamic network, all living organisms are related to each other (1). As the oldest and most widespread means of communication, chemical cues and signals are widely used between organisms (2). The chemical cues or signal transfer information between individuals of the same species are usually called pheromones (3) and are used in a multitude of contexts, including mate localization and mate choice as well as other social interactions in insects (2, 4). Most insect pheromones are thought to be synthesized from fatty acids, isoprenoids or amino acids in the glands of the insects (5). However, an increasing number of studies have revealed that microorganisms, including host-associated and environmental microorganisms, can also have an impact on chemical communication between individuals of the same insect species. To increase rates of transmission between host individuals, some parasites or pathogens of insects can manipulate the host’s behavior or physiology by affecting the host’s odor profiles (especially aggregation pheromones) and directly altering host communication with conspecifics (6, 7). In addition, the ecology of insects is influenced by microbial symbionts. Some microbial symbionts may influence the development of the host nervous system, including the photo- or chemosensory system (8). Reproductive manipulators such as *Wolbachia, Rickettsia, Cardinium, Spiroplasma, Arsenophonus* and some Bacteroidetes bacteria may also actively interfere with their host’s reproductive behavior to facilitate their own spread within the host population (2).

Volatile molecules produced by the microbiota play a primary role in chemical communication between insects (1), and direct production of pheromone components by the microbiota is one of the most obvious mechanisms (2). In this context, not only tightly insect-associated microbes but also many microbial fermentation products could be detected by insects (9). For some extracellular microbes in the secretions, excretions or secretory glands of insects, direct production of pheromones is of great importance (10). Since many pheromones are derived from some basic metabolites, such as fatty acids, amino acids, terpenoids and sterols (5, 11), the microbiota can easily regulate the production of pheromones by supplying or depleting precursor metabolites for pheromone synthesis (12). For example, the gut bacterium *Pantoea agglomerans* can produce the aggregation signal guaiacol in *Schistocerca gregaria* (13). In *Blattella germanica,* succinic acid and other acids used as aggregation pheromones are also associated with gut bacteria (14). In *Anopheles gambiae,* some bacteria can produce pyrazines, carboxylic acids, and alcohols and thereby stimulate oviposition (15). In eusocial insects, social behaviors could also be controlled by microbiota-produced pyrazines, cuticular hydrocarbons, and other semiochemicals (16). Though many insect pheromones were hypothesized to be of microbial origin due to their molecular structures that suggest biosynthetic pathways usually not present in higher eukaryotes, direct evidence for a microbial production remains lacking (2).

Tephritidae containing over 4,000 species, pose an enormous threat to fruit and vegetable production worldwide (17). Studying the reproductive behavior of agricultural pests could not only help elucidate the mechanism of mate selection but also provide useful information for improving control strategies (17). Among Tephritidae, males are widely believed to produce sex pheromones to attract females (18, 19). In *B. dorsalis,* studies have indicated that both sexes release molecules that attract conspecifics of the opposite sex. Baker and Bacon (1985) demonstrated that several spiroacetals in females could attract males (20). However, subsequent studies showed that males produce several volatiles in the rectal glands with pheromonal activity towards females (21). Moreover, other studies have extended the list of male-borne pheromones by adding other molecules (22). By summarizing the previous literature, we found that most sex pheromones in Tephritidae were identified in the rectal glands (17), which may indicate their relationship with the rectal microbiota. Here, we reinvestigated the sex pheromones in the rectal glands of *B. dorsalis.* We demonstrated that 2,3,5-trimethylpyrazine (TMP) and 2,3,5,6-tetramethylpyrazine (TTMP) are male-borne sex pheromones and proved that rectum symbionts are responsible for producing TMP and TTMP *in vitro.*

## Results

### Sex pheromones are produced in the rectums of mature males

We used 12-day-old unmated females in behavioral assays because they are highly attracted to sex pheromones produced by males. To test whether sex pheromones were present in the rectums of the males, the attraction effect of male rectum extracts was tested in a Y-tube olfactometer. For this purpose, 400 μl extracts were added as the attractants. The olfactometer arm containing extracts from the rectums of 9- and 12-day-old males attracted significantly more females than the control arm (Figure 1A). Anatomical observations of the male rectums at different stages showed that they were significantly enlarged and filled with fluid at the later stages (9- and 12-day-old males) (Figure 1B). These results indicate that male rectums at later stages may release volatiles that can attract females. To identify these attractants, extracts from the male rectums were analyzed with gas chromatography-mass spectrometry (GC-MS), and a marked difference was observed among stages; TMP, 2-ethyl-3,5-dimethylpyrazine, TTMP and 2,3,5-trimethyl-6-ethylpyrazine were present in the rectums at the later stages but absent at the early stages (Figure 1C and 1D). These results indicate that TMP, 2-ethyl-3,5-dimethylpyrazine, TTMP and 2,3,5-trimethyl-6-ethylpyrazine likely act as sex pheromones and are produced by mature males.

**Figure 1.**
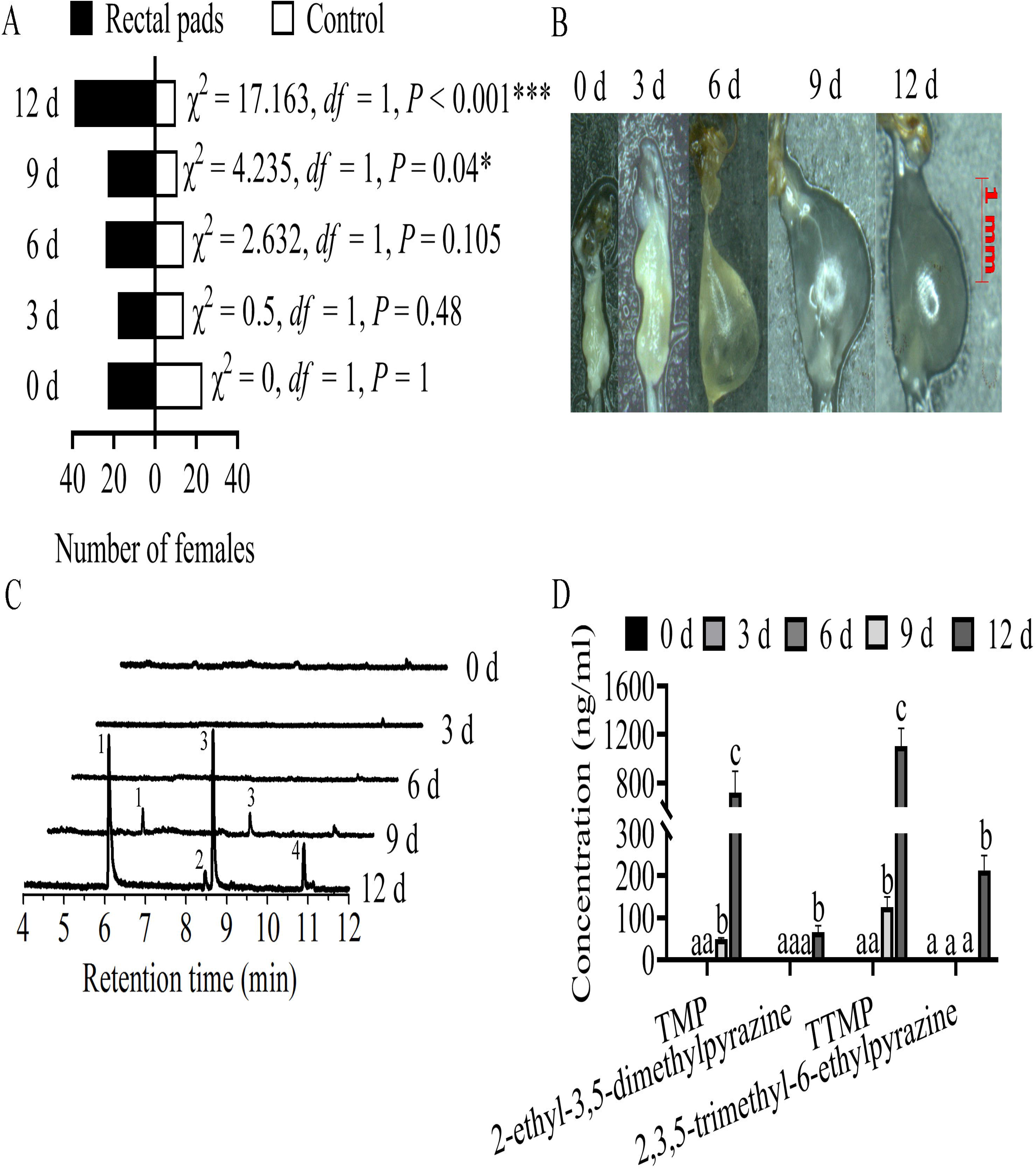
Attraction effects of male rectal extracts on mature virgin females and identification of chemicals in the extracts. (A) Attraction of mature virgin females by the extracts. (B) Morphology of the male rectum at different ages. (C) Chemical identification of rectal extracts. The numbers 1, 2, 3 and 4 represent TMP, 2-ethyl-3,5-dimethylpyrazine, TTMP and 2,3,5-trimethyl-6-ethylpyrazine, respectively. (D) Content comparison of TMP, 2-ethyl-3,5-dimethylpyrazine, TTMP and 2,3,5-trimethyl-6-ethylpyrazine in male rectums at different ages. Asterisks indicate significant differences (\**P* < 0.05, *** *P* < 0.001). Different letters indicate significant differences between groups (ANOVA with Tukey’s test).

### TMP and TTMP in the male rectum are sex pheromones

To explore candidate compounds in the male rectum that act as sex pheromones, we examined the electroantennography detection (EAD) signals of the extracts from rectums of 12-day-old males in the antennae of 12-day-old females. The results indicated that TMP and TTMP in the rectums could elicit an EAD response in females (Figure 2A). Y-tube olfactometer assays showed that 2000 ng/ml TMP and TTMP could significantly attract female flies (Figure 2B and 2C). Since TMP and TTMP were both detected in the extracts and their ratio was approximately 1:1, we compared the attraction effect of their mixture with that of TMP or TTMP. The results showed that the attraction effects on the females were similar for the mixture and TMP or TTMP when the concentration of the mixture was 1000 ng/ml (both TMP and TTMP were present at 1000 ng in the 1 ml mixture) (Figure 2D and 2F). Taking these results into account, we used mixtures of TMP and TTMP in subsequent experiments. In trap assays, we also found that mixtures of TMP and TTMP could attract significantly more females (Figure 2F). However, the effective concentration was 1 mg/ml, which was much higher than the concentration used in the Y-tube olfactometer assays.

**Figure 2.**
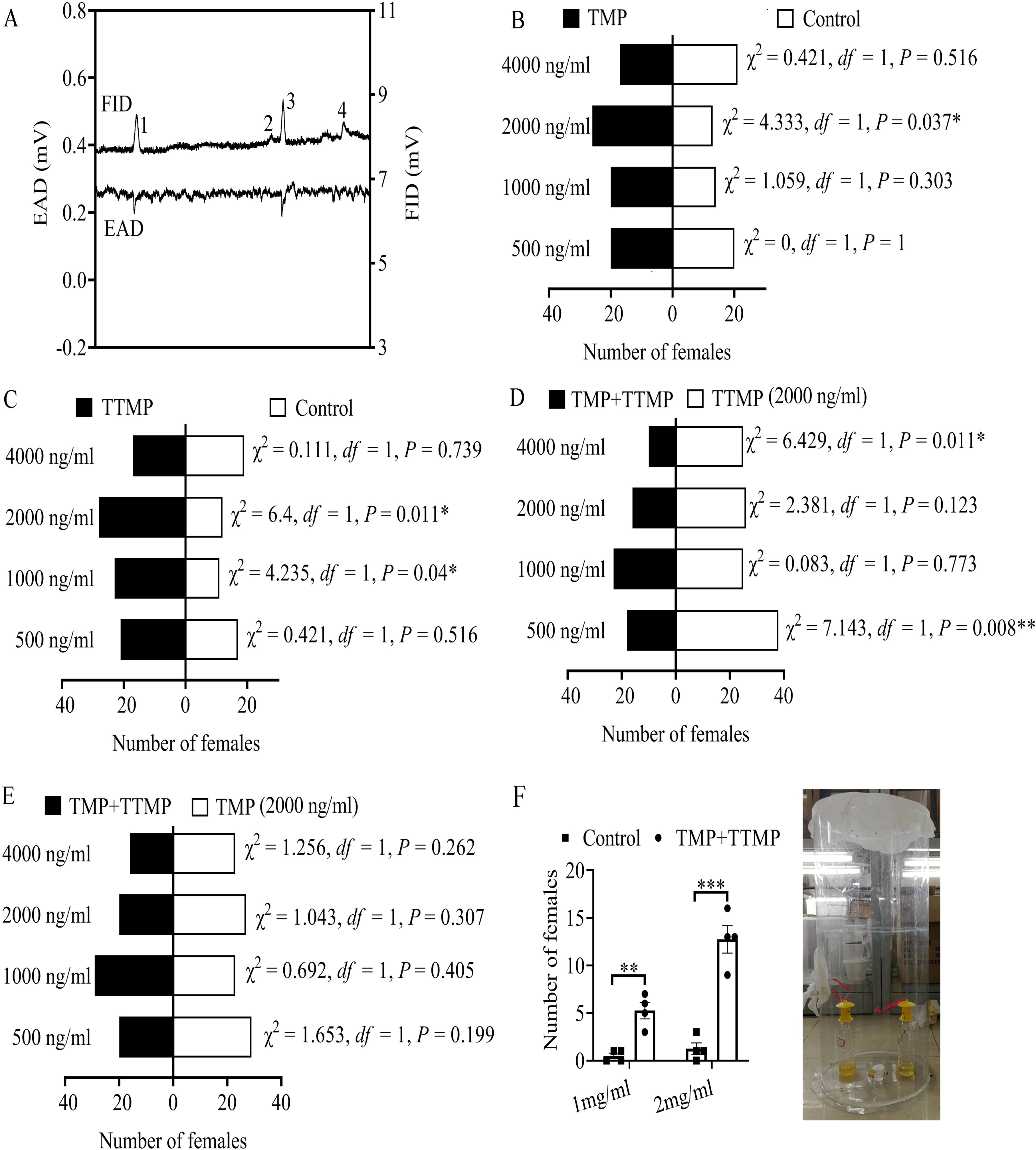
Attraction effects of TMP and TTMP on mature virgin females. (A) GC-EAD profile of rectal extracts of 12-day-old males in mature virgin females. The numbers 1, 2, 3 and 4 represent TMP, 2-ethyl-3,5-dimethylpyrazine, TTMP and 2,3,5-trimethyl-6-ethylpyrazine, respectively. (B) Attractiveness of TMP to mature virgin females. (C) Attractiveness of TTMP to mature virgin females. (D) Attraction effects on mature virgin females compared between a mixture of TMP and TTMP and TTMP alone. (E) Attraction effects on mature virgin females compared between a mixture of TMP and TTMP and TMP alone. (F) Capture effect of the mixture of TMP and TTMP on mature virgin females. In all figures, asterisks indicate significant differences (\**P* < 0.05, \*\**P* < 0.01).

### Rectal bacterial diversity changes significantly with the development of males

Previous studies have indicated that many bacteria in Bacilli have the pathway to produce TMP and TTMP (23–26). Moreover, no research had indicated that pyrazines could be synthesized by insects themselves till now. Thus, we hypothesized that rectal bacteria might be involved in producing TMP and TTMP. To test this hypothesis, we localized the bacteria present in the rectum of male flies and investigated bacterial diversity by 16S rRNA sequencing. Using the Cy3-labeled general eubacterial probe, fluorescence in situ hybridization (FISH) visualization was performed, and it revealed that many bacteria were present in the male rectum (Figure 3A). qPCR results showed that the total bacterial content in the male rectum increased significantly with development, especially in 6-day-old males (Figure 3B). 16S rRNA gene sequencing results indicated that the rectal bacteria communities in 6-, 9- and 12-day-old males differed significantly from 0- and 3-day-old males (Figure 3C), the dominant bacteria present in the rectums of newly emerged males belonged to Gammaproteobacteria, Alphaproteobacteria and Bacilli, while almost 80% of the total bacteria in 3-, 6-, 9- and 12-day-old males belonged to Bacilli (Figure 3D). By comparing the relative abundances of Bacilli at different times, Bacilli increased significantly with the development of the males (Figure 3E). Moreover, bacterial community profiles in the rectums revealed significant differences in operational taxonomic units (OTUs) and the Shannon diversity index over time (Figure S1). These results indicate that the changed rectal Bacilli may be associated with the sex pheromones generated in 9- and 12-day-old males.

**Figure 3.**
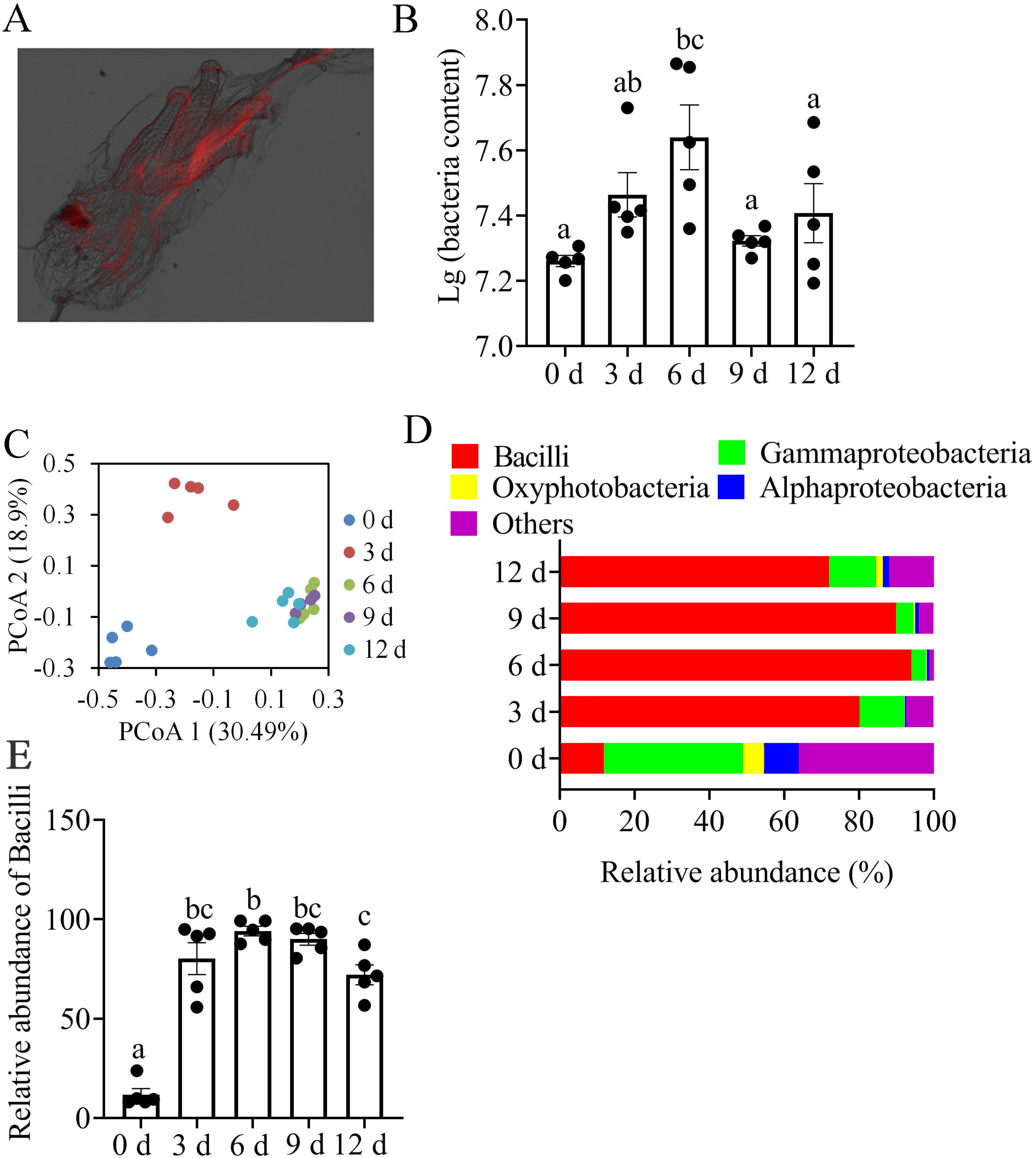
Rectal bacterial investigations. (A) Bacterial localization in the male rectum. Red signals indicate rectal bacteria. (B) Absolute number of bacteria in rectums of males at different ages. (C and D) Rectal bacterial composition identified by 16S rRNA sequencing. (F) Relative abundance of Bacilli in rectums of males at different ages. In all figures, different letters above the bars indicate significant differences between groups (ANOVA with Tukey’s test).

### Rectal bacteria contribute to sex pheromones

To test whether the sex pheromones were influenced by the bacteria associated with rectums, we fed male flies with antibiotics. Nine days later, qPCR results showed that the total bacterial content in rectums decreased significantly after feeding the flies with antibiotics (Figure 4A). The rectum (in 6-, 9- and 12-day-old males) sizes were decreased significantly after treatment with streptomycin and ceftriaxone, while fosfomycin treatment had no effect on rectum size (Figure 4B). Moreover, TMP and TTMP contents in rectums were decreased significantly after treatment with streptomycin and ceftriaxone, while fosfomycin treatment had no effect on TMP and TTMP contents (Figure 4C). 16S rRNA gene sequencing results indicated that the rectal bacteria communities in antibiotics treated males differed significantly from the control males (Figure 4D), the dominant bacteria present in control and fosfomycin-treated rectums belonged to Bacilli, while the abundance of Bacilli decreased significantly after treatment with streptomycin and ceftriaxone (Figure 4E and 4F). Moreover, bacterial community profiles in rectums at different times indicated that there were significant differences in OTUs and Shannon diversity indexes (Figure S2). These results indicated that the bacteria in Bacilli may be involved in producing TMP and TTMP.

**Figure 4.**
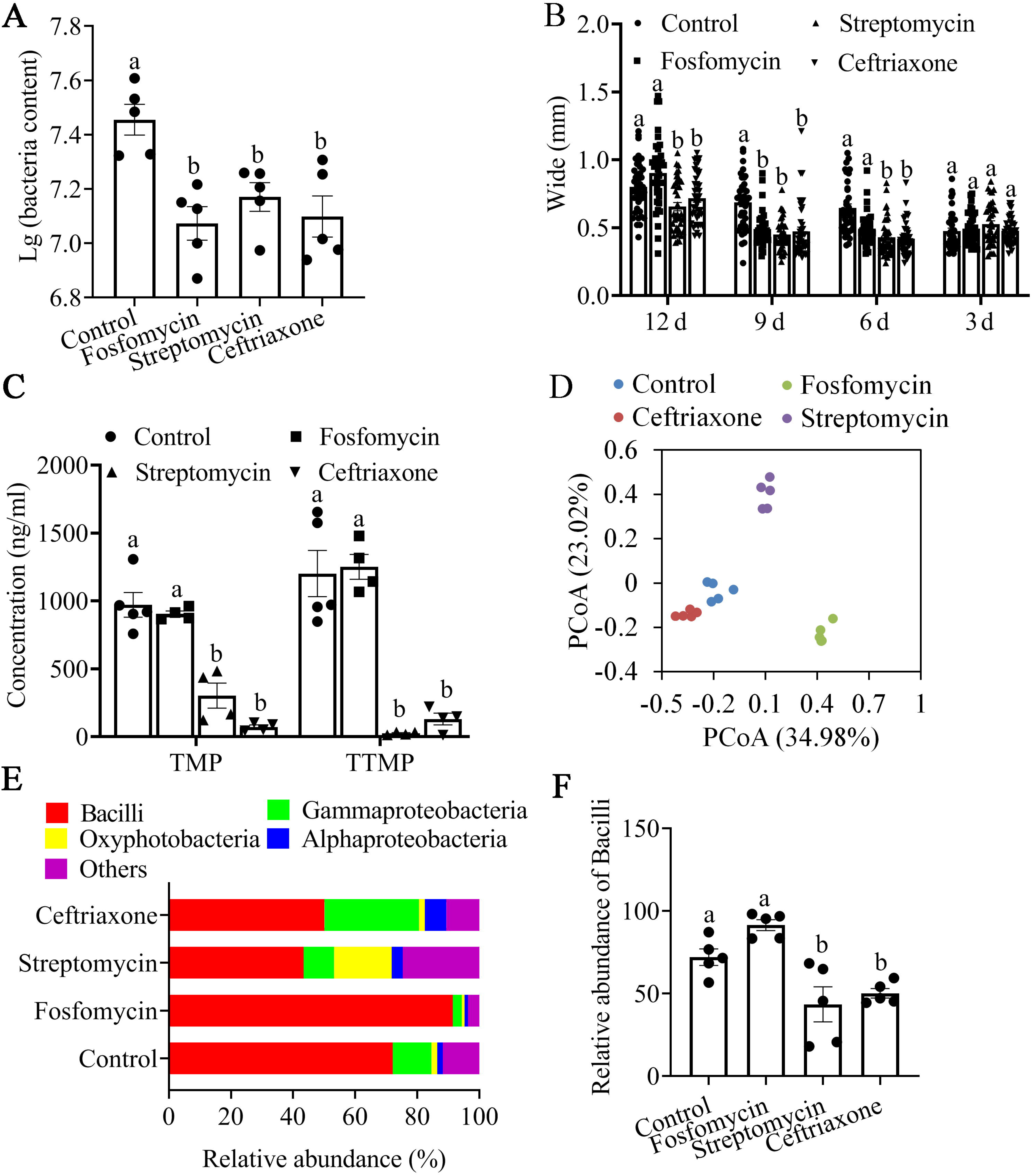
Effect of antibiotic treatments on rectal bacterial composition and volatile contents. (A) Effect of antibiotic treatments on the absolute number of bacteria in the rectums. (B) Effects of antibiotic treatments on rectum size. Different letters indicate significant differences between groups (ANOVA with Tukey’s test). (C) Effects of antibiotic treatments on TMP and TTMP contents in the rectum. (D and E) Rectal bacterial composition identified by 16S rRNA sequencing. (F) Relative abundance of Bacilli in rectums of males treated with different antibiotics. In all figures, different letters above the bars indicate significant differences between groups (ANOVA with Tukey’s test).

### Bacterial isolates from rectums produce TMP and TTMP *in vitro*

To verify whether the rectal bacteria could produce TMP and TTMP, the bacteria were isolated. As a result, four bacterial strains belonging to Bacilli were identified: *Lactococcus garvieae* (accession number: MW041150), *Bacillus pumilus* (accession number: MW041151), *Bacillus altitudinis* (accession number: MW041152) and *Bacillus safensis* (accession number: MW041153). Antibiotic sensitivity tests showed that all the isolated bacteria were sensitive to streptomycin and ceftriaxone, while they were not sensitive to fosfomycin (Figure 5A and 5B). These results were in accordance with those shown in Figure 4E and 4F. Moreover, *in vitro* fermentation assays indicated that *B. pumilus, B. altitudinis* and *B. safensis* had strong ability to produce TMP and TTMP, when glucose and threonine were supplied to the bacteria as substrates (Figure 5C). For TMP, the concentrations in the fermented broths of the isolated bacteria were higher than 2 μg/ml after 6 days (Figure 5D). For TTMP, the concentrations in the fermented broths were higher than 6.8 μg/ml (Figure 5D). Previous study had proposed the synthesis mechanisms of TMP and TTMP in *Bacillus* (26). In the proposed alkylpyrazine synthesis pathways, glucose and threonine are the key substrates for synthesizing TMP and TTMP (26). In the synthesis pathway, changes of glucose contents have significant influence on synthesis TMP and TTMP (26). Thus we have investigated the profiles of glucose and threonine (especially glucose) in the male rectums. The results indicated that both glucose and threonine could be identified in rectums of the male flies (Figure 5E and 5G). Moreover, the contents of the key substrate-glucose were significantly higher in rectums of 9- and 12- day-old males (Figure 5F), which was consistent with higher TMP and TTMP contents in 9- and 12- day-old males. These results indicated TMP and TTMP in the rectums of *B. dorsalis* could be produced by *Bacillus* in rectums using glucose and threonine as substrates and the increased contents of glucose may be the factors that affect TMP and TTMP contents in rectums of the male flies.

**Figure 5.**
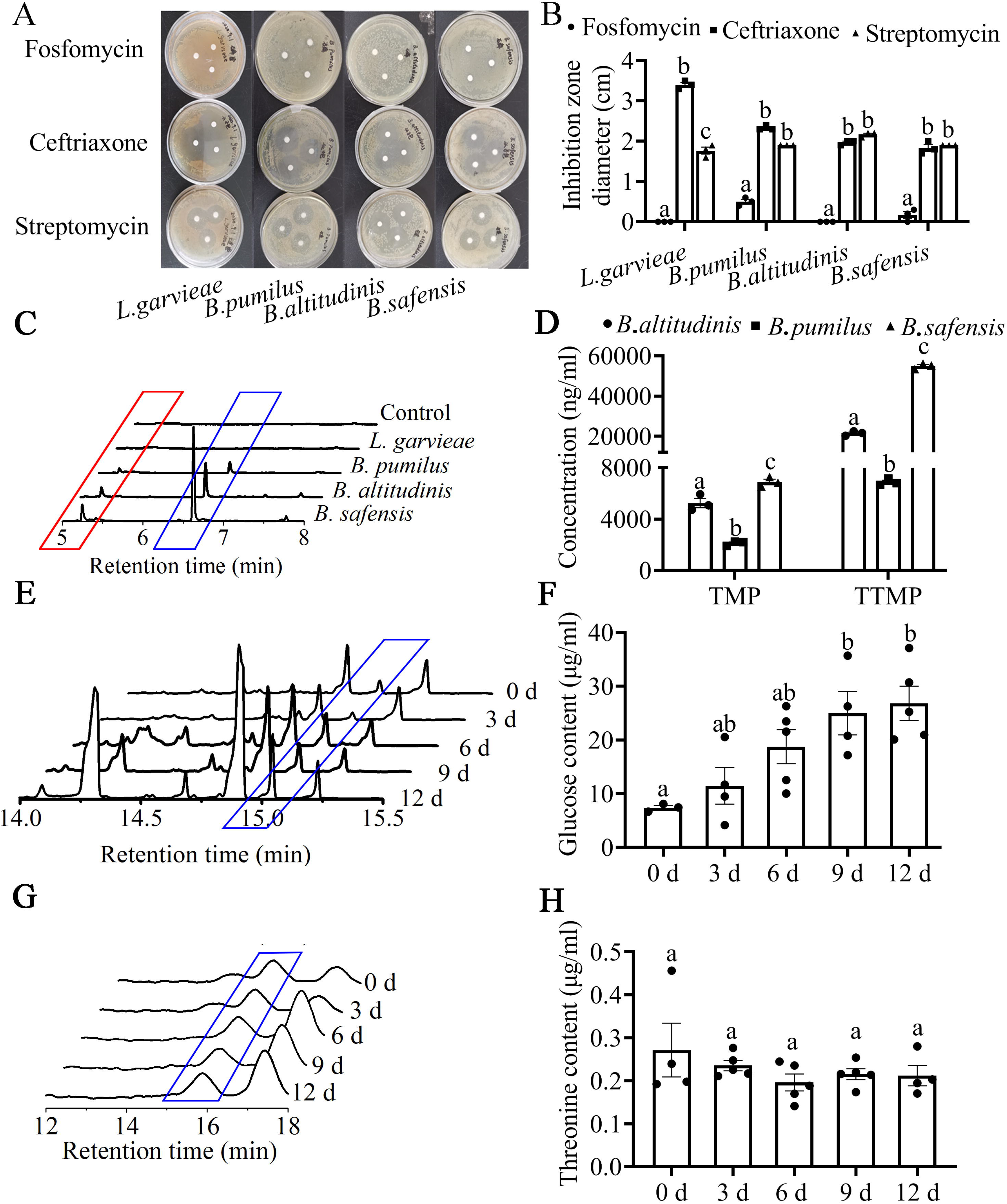
Isolation of rectal bacteria and their abilities to produce TMP and TTMP. (A and B) Sensitivity of isolated bacteria to antibiotics. (C) TMP and TTMP identification in the fermented broth of the isolated bacteria. The peaks in the red and blue boxes represent the gas chromatogram traces of TMP and TTMP, respectively. (D) TMP and TTMP concentrations in the fermented broths of the isolated bacteria. (E) Glucose identification in the rectums of the male flies. The peaks in the blue box represent the gas chromatogram traces of glucose. (F) Glucose concentrations in rectums of males at different ages. (G) Threonine identification in the rectums of the male flies. The peaks in the blue box represent the liquid chromatogram traces of threonine. (F) Threonine concentrations in rectums of males at different ages. In all figures, different letters above the bars indicate significant differences between groups (ANOVA with Tukey’s test).

## Discussion

In tephritid flies, courtship and mating can be influenced by olfactory cues, and olfactory stimuli usually play a key role in the search for mates (17). Although studies have indicated that tephritid males produce sex pheromones to attract females (27), both sexes of *B. dorsalis* produce sex pheromones (19, 20). Perkins et al. (1990) (22) claimed that the list of male-borne pheromones includes TMP in the rectum. By reinvestigating the volatile profiles in the rectums of males, we identified both TMP and TTMP for the first time. We found that TMP and TTMP were present only in the rectums of mature males. By testing their functions, we found that both molecules showed selective attraction towards mature females. Moreover, both TMP and TTMP have be identified in other tephritid flies (28), which may indicate their shared origin.

Researchers have hypothesized that microbes influence aggregation, sexual communication and mate choice in insects since experimental antibiotic disruption of host-associated microbial communities results in reduced attractiveness and fecundity (14, 29, 30). Specifically, dominant gut bacteria, especially *Klebsiella* and *Enterobacter,* have been reported to influence the mating behavior of tephritid fruit flies (31–33). However, it remains unclear whether these effects are mediated by a direct bacterial contribution to the flies’ chemical profiles or by bacterium-provided benefits that enhance the fitness of the flies and thereby improve their pheromone composition or overall physical competitiveness. To date, the only conclusive case of direct microbial production of a sex pheromone was reported by Hoyt et al. (1971) (34). This study revealed that the female sex pheromone phenol 1 in the New Zealand grass grub beetle *Costelytra zealandica* is produced by a bacterium in the colleterial glands. Although some other insect pheromones are also hypothesized to be of microbial origin (35), direct evidence for microbial production remains scarce. In our study, we demonstrated that bacteria in the rectum of *B. dorsalis* can produce sex pheromones *in vitro,* providing another example of direct microbial production of sex pheromones. Such a mechanism may be widespread in tephritid fruit flies since many tephritid fruit flies use rectally produced TMP and TTMP as sex pheromones.

Pyrazines, including TMP and TTMP, are an important class of bacterial volatile organic compounds with chemical signaling properties in insects (36). Although many studies have indicated that pyrazines can be biosynthesized by bacteria, only a few have shown their possible origin from insect symbionts. In our study, we demonstrated that the symbionts in the rectum of *B. dorsalis* can produce TMP and TTMP, which can significantly attract female flies. TMP and TTMP can be produced by bacteria of the genus *Bacillus* during industrial fermentation (26). The high abundance of *Bacillus* in the rectum of *B. dorsalis* may also indicate their important role as symbionts in the lifecycle of *B. dorsalis.* In recent decades, studies have demonstrated the important role of pyrazines in eliciting the behavior (e.g., acting as an attraction, alarm, and trailing pheromone) of insects (37–39), and these pyrazines are usually isolated from the glands of insects (40–42). These glands, which are moist, nutrient-rich, and largely anaerobic, should be conducive to the proliferation of symbiotic, particularly fermentative, bacteria. Thus, the fermentation hypothesis for insect chemical communication posits that as bacteria ferment or otherwise metabolize the nutrient-rich substrates in the glands, they generate odorous metabolites that are subsequently used by their hosts to communicate with conspecifics (43). As a supplement to our study, investigations should be performed to reveal whether the pyrazines used as pheromones in the glands of insects (such as *Solenopsis invicta, Pachycondyla sennaarensis* and *Wasmannia auropunctata* (Roger)) are produced by symbionts in the glands.

This study provides strong empirical support for the fermentation hypothesis of chemical communication in *B. dorsalis.* However, only three strains that could produce sex pheromones were isolated from the rectum, and antibiotic-fed males did not completely lose TMP and TTMP in the rectum. Therefore, we cannot exclude the possibility that some other bacteria can also produce TMP and TTMP. It is also possible that TMP and TTMP are produced by the fly itself. Further tests could be performed to investigate whether the genomes of the isolated bacteria contain genes coding for the fermentation pathways leading to the production of TMP and TTMP.

## Materials and methods

### Flies used in this study

All experiments were carried out with a laboratory-reared *B. dorsalis* strain collected from a carambola *(Averrhoa carambola)* orchard located in Guangzhou, Guangdong Province, in April 2017. The flies were maintained in the laboratory under the following conditions: 16:8 h light:dark cycle, 70-80% relative humidity (RH), and 25 ± 1°C. A maize-based diet containing 150 g corn flour, 150 g banana, 0.6 g sodium benzoate, 30 g yeast, 30 g sucrose, 30 g paper towel, 1.2 ml hydrochloric acid and 300 ml water was used to feed the larvae, and the adult diet consisted of water (with or without added antibiotics), yeast hydrolysate, and sugar. Male and female flies were reared separately upon emergence. For antibiotic treatments, newly emergent male flies were fed fosfomycin (3 mg/ml, diluted in sterile water), streptomycin (3 mg/ml, diluted in sterile water) or ceftriaxone (3 mg/ml, diluted in sterile water). The flies were continuously fed the antibiotics.

### Olfactometer bioassays

An olfactometer consisting of a Y-shaped glass tube with a main arm (20 cm length*5 cm diameter) and two lateral arms (20 cm length, 5 cm diameter) was used. The lateral arms were connected to glass chambers (20 cm diameter, 45 cm height) in which the odor sources were placed. To ensure a supply of odor-free air, both arms of the olfactometer received charcoal-purified and humidified air at a rate of 1.3 L/min.

To test the attraction effect of male rectum extracts on females, the rectums of 60 males (0, 3, 6, 9 and 12 days old) were collected and extracted with 400 μl ethanol for 24 h. The prepared extract was placed in one odor glass chamber. In the control odor glass chamber, 400 μl ethanol was placed. Then, 12-day-old unmated females were individually released at the base of the olfactometer and allowed 5 min to show a selective response. The response was recorded when a female moved more than 3 cm into one arm and stayed for >1 min. Females that did not leave the base of the olfactometer were recorded as nonresponders. Only females that responded were included in the data analysis. Odor sources were randomly placed in one arm or the other at the beginning of the bioassay, and the experiment was repeated ten times. The system was washed with alcohol after every experiment. More than 30 females were selected for testing, and each female was used only once for each odor.

To test the attraction effect of TMP and TTMP on females, 1 ml TMP or TTMP at different concentrations (diluted in ethanol) was placed in one odor glass chamber. In the control odor glass chamber, 1 ml ethanol was placed. Then, the selective behavior of 12-day-old unmated females was tested as described above. Odor sources were randomly placed in one arm or the other at the beginning of the bioassay. After 10 replicates, the system was washed with alcohol. For each odor, more than 30 females were selected for testing, and each female was used only once.

To test the attraction effect of the mixture of TMP and TTMP on females, a 1 ml mixture of TMP and TTMP (4000, 2000, 1000 and 500 ng/ml diluted in ethanol, both TMP and TTMP at 4000, 2000, 1000 or 500 ng in 1 ml ethanol) was placed in one odor glass chamber. In the control odor glass chamber, 1 ml TMP or TTMP solution (2000 ng/ml) was placed. Then, the selective behavior of 12-day-old unmated females was tested as described above. Odor sources were randomly placed in one arm or the other at the beginning of the bioassay. After 10 replicates, the system was washed with alcohol. For each odor, more than 30 females were selected for testing, and each female was used only once.

### Volatile analysis

The volatiles present in the extracts prepared as described above were identified by GC-MS. Compounds were analyzed by GC-MS with an Agilent 7890B Series GC system coupled to a quadrupole-type-mass-selective detector (Agilent 5977B; transfer line temperature: 230°C, source temperature: 230°C, ionization potential: 70 eV). The volatiles were chromatographed on an HP-5 MS column (30 m, 0.25 mm internal diameter, 0.25 μm film thickness). Helium at a constant pressure of 110.9 kPa was used for carrier gas flow. One microliter extract was injected into the injector port (240°C). The column temperature was maintained at 50°C for 1 min, increased to 140°C at 5°C/min, increased to 250°C at 10°C/min and maintained for 3 min. Volatile identification was based on the comparison of mass spectra with those listed in the NIST mass spectral library. Additionally, the identification of TMP and TTMP was confirmed by comparing their retention times and mass spectra with those of authentic standards purchased from suppliers. Experiments were conducted in triplicate. To determine the contents of TMP and TTMP in extracts, a standard curve was generated with the authentic standards. The standard curves were prepared in triplicate (n=3) using 100, 10, 1 and 0.1 μg/ml solutions of TMP or TTMP (diluted in ethanol).

To identify the volatiles in the fermented broth of bacteria isolated from the male rectum, a 100-μm polydimethylsiloxane (PDMS) SPME fiber (Supelco) was used to capture the volatiles for 30 min at room temperature. Compounds were analyzed by GC-MS with an Agilent 7890B Series GC system coupled to a quadrupole-type-mass-selective detector (Agilent 5977B; transfer line temperature: 230°C, source temperature: 230°C, ionization potential: 70 eV). The fiber was inserted manually into the injector port (240°C) for desorption, and the volatiles were chromatographed on an HP-5 MS column (30 m, 0.25 mm internal diameter, 0.25 μm film thickness). Helium at a constant pressure of 110.9 kPa was used for carrier gas flow. After fiber insertion, the column temperature was maintained at 50°C for 1 min, increased to 140°C at 10°C/min, increased to 250°C at 25°C/min and maintained for 3 min. The fermented broth was prepared by adding isolated bacteria into 100 ml Luria-Bertani (LB) liquid medium supplemented with membrane filtration-sterilized threonine (42 mM), glucose (56 mM), and (NH_4_)2HPO_4_ (23 mM). Then, the bacteria were cultured aerobically at 37°C by shaking at 200 rpm for 6 days. Experiments were conducted in triplicate.

### GC-EAD analysis

GC-EAD analysis was performed to determine whether TMP and TTMP could elicit antennal responses in unmated mature females. The gas chromatograph (Agilent 7890A, USA) was equipped with an HP-5 capillary column (30 m ☐×☐ 0.32 mm ☐×☐ 0.25 μm, Agilent). The column temperature was maintained at 50°C for 1 min, increased by 5°C/min to 140°C, increased by 10°C/min to 250°C, and maintained for 10 min. Nitrogen was used as the carrier gas at 25 ml/min. One microliter extract was injected into the GC column (splitless model) at 250°C with a flame ionization detector (FID) at 260°C. For EAD preparations, an antenna of a female was mounted between two microelectrodes with electrode gels. The antenna tip was cut slightly to facilitate electrical contact. Only one recording was made per antenna, with ten successful recordings performed in total. The signals from the antennae were analyzed with GC-EAD 2014 software (version 4.6, Syntech).

### Trap assays

We adapted the assay from Figure 2F to compare attraction to a mixture of TMP and TTMP and fermented broth of the isolated bacteria. The test chamber was assembled with a plastic cylinder (120 x 30 cm) covered by a ventilated lid. The test chamber contained an odor-baited trap and a control trap. The traps were made of transparent plastic vials (20 x 6 cm) and were sealed with a yellow lid on which small entrances were set to let the flies in. To test attraction to a mixture of TMP and TTMP, 30 12-day-old unmated females were used for each trial. The odor-baited trap contained 50 ml TMP and TTMP mixture diluted in ethanol. The control trap contained an equal volume of ethanol. The experiment lasted for 24 h.

### Observation of rectal bacteria with FISH

The rectum of a 12-day-old male was dissected. The hybridization protocol for the rectum was approximately according to Cheng’s (44) methods. Briefly, the rectum was collected and immediately soaked in Carnoy’s fixative for 12 h. After sample fixation, proteinase K (2 mg/mL) treatment for 20 min at 37°C and HCl (0.2 mol/L) treatment for 15 min at room temperature were performed successively. Then, followed by dehydration in ethyl alcohol, the samples were incubated in buffer (20 mM Tris-HCl (pH 8.0), 0.9 M NaCl, 0.01% sodium dodecyl sulfate, 30% formamide) containing 50 nM Cy3-labeled general eubacteria probe EUB338 (5′-GCTGCCTCCCGTAGGAGT-3′) (45) for 90 min. After incubation, the samples were washed with buffer (0.1 M NaCl, 20 mM Tris/HCl (pH 8.0), 5 mM ethylenediaminetetraacetic acid (pH 8.0), 0.01% SDS) and observed under an epifluorescence microscope (AxioPhot, Carl Zeiss, Shinjuku-ku, Japan).

### Rectal bacterial identification

To analyze male rectal bacterial diversity, 5 male rectums were collected from 0-, 3-, 6-, 9- and 12-day-old males (five replicate samples were prepared for each time point). Moreover, 5 male rectums were also collected from the antibiotic-fed males (9 days old) (five replicate samples were prepared for each fly strain). Then, the bacterial DNA in the rectum was extracted using the Bacterial Genomic DNA Extraction Kit (Tiangen, Beijing, China, http://www.tiangen.com/asset/imsupload/up0250002001571219042.pdf) according to the manufacturer’s protocols. Before sequencing, qPCR was used to estimate the absolute abundance of bacteria in the rectum according to the method of Li et al. (2020) (46). To analyze the bacterial diversity in the male rectum at different ages and with or without antibiotic treatment, the 16S rRNA V3-V4 region was amplified with PCR according to the method of Li et al. (2020) (46). Then, the amplicons were purified and sequenced (2 × 250) on an Illumina HiSeq 2500 platform; the sequenced raw reads for each sample were cleaned and analyzed according to the standards of Li et al. (2020) (46). Sequencing data were submitted to NCBI, accession number: PRJNA669205.

For male rectal bacterial isolation, 5 rectums of 12-day-old unmated males were dissected and collected in a sterile centrifuge tube, to which 1 ml sterile water was added. The rectums were then ground with sterile grinding pestles and shaken for 20 min. A 200 μL volume of the diluted liquid was then used to coat an LB plate and cultured for 1 day. Pure bacterial colonies were selected by subculturing on LB media and stored in a 25% glycerol solution at −80°C. A Bacterial Genomic DNA Extraction Kit (Tiangen, Beijing, China) was used to extract bacterial DNA according to the manufacturer’s instructions. Universal primers (F: 5′-AGAGTTTCATCCTGGCTCAG-3′ and R: 5′-TACGGTTAXXTTGTTACGACTT-3′) were used to amplify the 16S rRNA. The PCR products were confirmed by electrophoresis on a 0.8% agarose gel, and the target PCR product was sequenced. The 16S rRNA sequence was BLAST searched against the NCBI NR database (https://blast.ncbi.nlm.nih.gov/Blast.cgi).

### Antibiotic testing of the isolated bacteria

To test bacterial sensitivities to antibiotics, 100 μl inoculum of the isolated bacteria was streaked onto LB agar flat plates, to which drug-susceptibility test papers were subsequently attached. On the papers, 5 μl fosfomycin (2 mg/ml), ceftriaxone (2 mg/ml) and streptomycin (2 mg/ml) were added. Then, the plates were incubated for 24 h at 37°C, and the diameters of the inhibition zones were measured.

### Glucose and threonine identification in male rectum

GC-MS analysis was done to identify glucose and threonine in rectums of males at different development stages. For glucose identification, sample preparation for GC-MS analysis was done similar to Mayack et al., (2020)(47). Briefly, 15 rectums of 0-, 3-, 6-, 9- and 12-day-old males were collected and put into a 1.5 ml microcentrifuge tube added 500 μl millipore sterile water, respectively. Then grind the sample with a grinding machine. The supernatant was collected by centrifugation into another centrifuge tube. To desiccate the samples, the samples were lyophilized overnight in a Lyophilizer Freeze Dryer (Labconco, Kansas City, MO, USA). Samples were then stored up to three days at −80 °C. The samples were derivatized to be prepared for GC-MS analysis. In each 1.5 ml microcentrifuge tube, 10 μl of pyridine and 20 μl of N, O-Bistrifluoroacetamide (BSTFA) with 1% Trimethylchlorosilane (TMCS) (Thermo Scientific, Waltham, MA, USA), were added. After thorough vortexing, samples were transferred to a 2 ml amber microvials with a 250 μl glass insert (Agilent Technologies, Santa Clara, USA). The samples were heated for 3 h at 70 °C and then stored at −20 °C until GC-MS analysis. For each run, 4 solutions of glucose ranging from 2.5 to 20 μg/ml were prepared to generate the standard curve for glucose quantification. These solutions were diluted, lyophilized, and derivatized as described above for the rectum samples. Samples and glucose standards were analyzed by Agilent 7890(GC)/5975C (MS). Injector temperature was maintained at 280 °C and the mass spectrometry source temperature at 230 °C. A DB5-MS capillary column was used. The injection temperature was held constant for 2 min at 65 °C. Temperature was increased by 10 °C per minute up to 140 °C, then increased to 250 °C at 25°C/min and maintained for 3 min. Helium was used as the carrier gas at a constant flow rate of 1.3 ml/min. The mass spectrometer was set to the electron impact mode and its ionization was set to 70 eV The scan mode was set at 1.27 scans per second, ranging from 50 to 650 DA, and samples were analyzed with a split time of 30 s. The typical retention times (~15 min) and prominent ion numbers (204.2) of glucose were used to identify glucose. Standard curves were generated from the peak areas with known glucose concentrations. For each rectum sample the standard curve linear equation was used to convert peak area into μg/ml concentrations.

For threonine identification, sample preparation for free amino acid analysis was done similar to Shahzad et al., (2019) (48). Briefly, 15 rectums of 0-, 3-, 6-, 9- and 12-day-old males were collected and put into a 1.5 ml microcentrifuge tube added 500 μl sulfosalicylic acid solution (5%, dilute in water), respectively. Then grind the sample with a grinding machine. The samples were centrifuged for 15 min with 12000 rpm. Then the supernatants were collected into another centrifuge tube and added 1ml more sulfosalicylic acid solution. Then threonine quantification for the samples was carried out by amino acid analyzer (HITACHI L-8900, Japan) with the standard method.

### Statistical analysis

A chi-square test was used to analyze the selective preference of females in the Y-shaped olfactometer. The results for trap assays were analyzed using independent-sample t-tests. Concentrations of the volatiles identified in the rectum of males of different ages, bacterial contents, abundances of Bacilli, rectal sizes and concentrations of glucose and threonine were compared with one-way analysis of variance (ANOVA) followed by Tukey’s post hoc tests. All statistical analyses were performed in SPSS 19.0.

## Acknowledgements

We are grateful to Guangzhou Genedenovo Biotechnology Co., Ltd for assisting in sequencing. The work was supported by the Natural Science Foundation of Guangdong Province (2019A1515012191).

## Notes

### Competing Interest Statement

The authors have declared no competing interest.

