## Supplemental figures for "Rectal bacteria produce sex pheromones in the male oriental fruit fly"

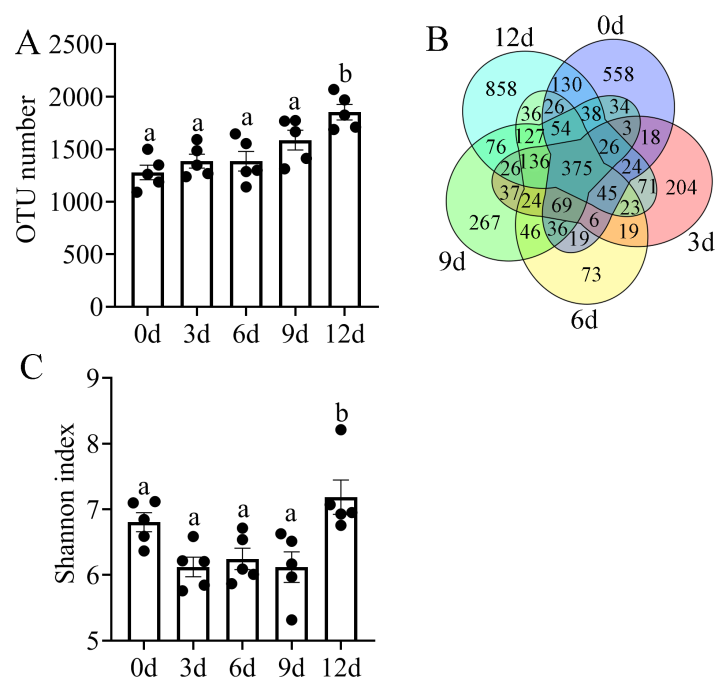

**Figure S1 Bacterial diversity in the rectums of males at different stages.** (A) OTU number of the bacteria. (B) Venn diagram showing the difference in OTUs between samples. (C) Shannon diversity index values of the bacteria. Different letters indicate significant differences in treatments detected by ANOVA at the 0.05 level.

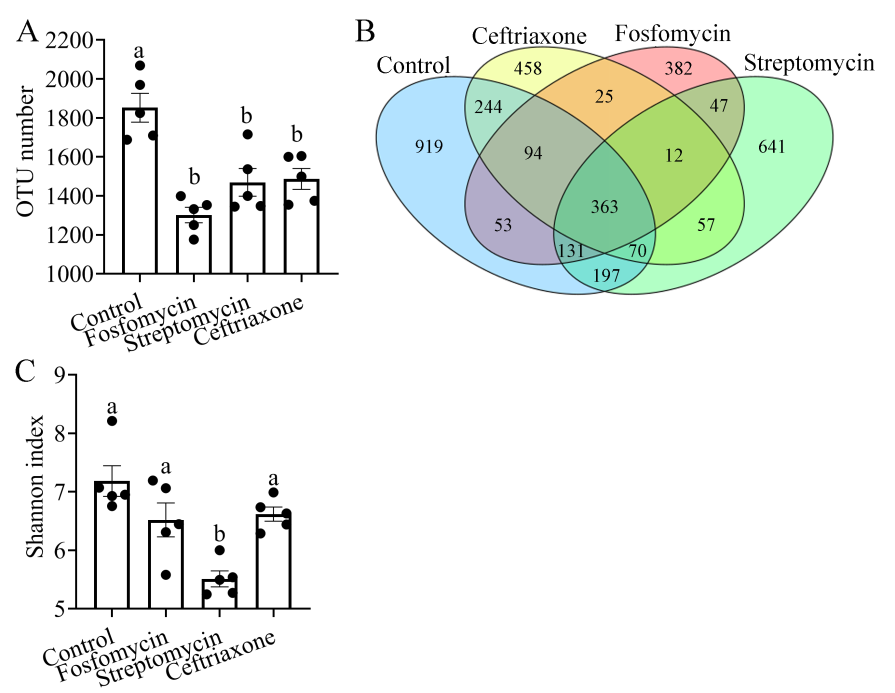

**Figure S2 Bacterial diversity in rectums of males treated with antibiotics.** (A) OTU number of the bacteria. (B) Venn diagram showing the difference in OTUs between samples. (C) Shannon diversity index values of the bacteria. Different letters indicate significant differences in treatments detected by ANOVA at the 0.05 level.
